# Evaluation of the Angus ICD9-CM Sepsis Abstraction Criteria

**DOI:** 10.1101/124289

**Authors:** Steven Horng, Larry A. Nathanson, David A. Sontag, Nathan I. Shapiro

## Abstract

**Objective:** Validate the infection component of the Angus International Classification of Diseases, Ninth Revision, Clinical Modification (ICD9-CM) sepsis abstraction criteria

**Design:** Observational cohort study

**Setting:** 55,000 visits/year Adult Emergency Department (ED)

**Patients:** All consecutive ED patient visits between 12/16/2011 and 08/13/2012 were included in the study. Patients were excluded if there was a missing outcome measure.

**Interventions:** None.

**Measurements and Main Results:** The primary outcome measure was suspected infection at conclusion of the ED work-up as judged by the physician. There were 34,796 patients who presented to the ED between 12/16/11 and 8/13/12, of which 31,755 (91%) patients were included and analyzed. The original Angus sepsis abstraction criteria had a sensitivity of 55%, specificity of 97%, PPV of 82%, NPV of 88%, accuracy of 87%, and a F_1_ score of 0.66. The modified Angus sepsis abstraction criteria which includes codes added after the original publication had a sensitivity of 65%, specificity of 96%, PPV of 81%, NPV of 91%, accuracy of 89%, and F_1_ score of 0.72.

**Conclusions:** In our study, the Angus abstraction criteria have high specificity (97%), but moderate sensitivity (55%) in identifying patients with suspected infection as defined by physician at the time of disposition from the emergency department. Given these findings, it is likely that we are underestimating the true incidence of sepsis in the United States and worldwide.

## Introduction

### Background

In 2001, Angus et. al, published an estimate of the incidence of severe sepsis in the United States using a set of ICD9-CM codes (1). This estimate of severe sepsis is the most commonly cited statistic for sepsis with 498 citations in PubMed and 2,674 citations in Scopus. This estimate uses ICD9-CM sepsis abstraction criteria consisting of two components: one set of ICD9-CM codes to identify infection and a second set of codes to identify acute organ dysfunction. Severe sepsis is then retrospectively identified by finding patients with both an ICD9-CM code for infection and a code for acute organ dysfunction. This methodology allows researchers to identify large cohorts of patients with severe sepsis for both epidemiological surveillance and cohort selection. A limitation to this method is that it does not utilize any additional clinical information from the medical records or clinicians themselves to assess the presence of severe sepsis.

### Importance

Septic shock is responsible for significant morbidity, mortality, and cost to patients in our healthcare system. Sepsis accounts for 1 in 5 admissions to the ICU and remains the leading cause of death in non-cardiac ICU’s. The hospital case mortality rate for severe sepsis is reported to be between 30-50% (2-5), leading to an estimated 751,000 deaths nationally (5). The overall cost of care is estimated at $16.7 billion annually in the US (5). Accurate statistics about the scope and magnitude of sepsis is essential to ensure that appropriate clinical, administrative, and research resources are allocated to combat this important disease process.

### Goals of This Investigation

Although the Angus abstraction criteria are widely used for epidemiological surveillance, prospective clinical validation of the criteria are limited. It is possible that the criteria could either be underestimating or overestimating the true incidence of severe sepsis. Although the use of ICD9-CM codes to identify infection has face validity, we wish to compare this to the assessment of ED providers when delivering clinical care.

The goal of this investigation is to prospectively validate the infection component of the Angus ICD9-CM sepsis abstraction criteria. Additionally, since many additional ICD9-CM codes consistent with infection have been added in the decade since the original publication of the Angus article, we have also developed an updated version of the Angus criteria to include these new ICD9-CM codes. In this study, we first validate the original Angus criteria and then evaluate a modified version.

## Materials and Methods

### Study Design

We conducted a prospective, observational cohort study of 31,755 consecutive patients at a 55,000 visits/year Emergency Department over a 242-day period. At the conclusion of the ED work-up, we asked the treating physician to determine whether they suspected the patient to have an infection. We then used the Angus ICD9-CM abstraction criteria to classify the patient as infected or not. We then compared the Angus ICD9-CM determination to the clinical assessment using the question asked at the time of ED disposition as the criterion standard. This study was approved by our Institutional Review Board.

### Setting and Selection of Participants

The study was performed in a 55,000 visits/year Level I trauma center and tertiary care academic teaching hospital. All consecutive ED patient visits between 12/16/2011 and 08/13/2012 were included in the study. Visits were excluded if the outcome measure of suspected infection was not answered by the primary treating physician in the emergency department.

### Outcome Measures

The primary outcome measure was suspected infection at conclusion of the ED work-up as judged by the physician. Although we could have used positive urine culture, chest radiograph, or other diagnostic test as the criterion standard for infection, we specifically chose suspected infection because the American College of Chest Physicians/Society of Critical Care Medicine diagnostic criteria (6) requires only suspected infection for a diagnosis of sepsis.

### Methods of Measurement

In order to admit, discharge, transfer, or otherwise disposition a patient from the ED, the Attending or Resident Physician caring for the patient must enter an order in our ED information system. As part of that disposition module, resident and attending physicians answered the voluntary question “Do you think this patient has an infection” and responds using a 5-point Likert Scale [1 – Unlikely, 3 – Unsure, 5 – Likely] and served as the primary reviewer. We also had one of three attending emergency physicians answer the same question while working clinically in the emergency department to serve as a second reviewer in order to measure inter-rater reliability. There was no overlap between the primary and secondary reviewers. The answer choices were then dichotomized, with choices 4 and 5 yielding a positive outcome measure of suspected infection.

### Abstraction Criteria

As part of our coding and billing process, a set of 10 or fewer ICD9-CM codes are assigned to each ED visit by trained coders based on physician documentation. These codes are generally assigned within 7 days of the ED visit. Although in-hospital ICD9-CM codes are also subsequently assigned for patients admitted to the hospital, we are specifically interested in the suspicion of infection in the emergency department, not infection that might develop or be diagnosed later in a patient’s hospital stay.

Since many additional ICD9-CM codes were added in the decade since the original Angus abstraction criteria were published, we manually reviewed the original criteria a priori. We identified several new ICD9-CM codes indicative of infection including codes for Viral Meningitis, Cholangitis, Orbital Cellulitis, as well as codes for Sepsis, Severe Sepsis, and Septic Shock. We report the additional codes in our modified Angus ICD9-CM Sepsis Abstraction Criteria and are reported in **Appendix 1**. The full listing of both the original and modified Angus abstraction codes are provided online as supplementary material in tab-delimited format.

### Primary Data Analysis

Means with 95% confidence intervals were reported for age. Counts with percentages were reported for gender, Emergency Severity Index, and disposition. Sensitivity, specificity, positive predictive value (PPV), negative predictive value (NPV), and accuracy were reported for both the original and the modified Angus abstraction criteria. An F_1_ score is also reported as a measure of test performance that takes into account the tradeoff between sensitivity and specificity, and represents the harmonic mean of sensitivity and positive predictive value 2 x (sensitivity x positive predictive value) / (sensitivity + positive predictive value). Finally, a sensitivity analysis was performed adding the top 2 ICD9-CM codes (nausea, vomiting, and diarrhea) associated with false negatives to the modified abstraction criteria. A sensitivity analysis was also performed by deleting the ICD9-CM code (fever) associated with false positives. Finally, we calculated a Cohen’s Kappa statistic between the primary and secondary reviewer as a measure of inter-rater reliability. Statistical analysis was performed using JMP (JMP, Version 9. SAS Institute Inc., Cary, NC, 1989-2010.)

## Results

There were 34,796 patients who presented to the ED between 12/16/11 and 8/13/12, of which 31,755 (91%) patients were included and analyzed. (**Figure 1)**. During this study, 263 physicians served as primary reviewers, providing outcome measure for a median of 50 patients (Interquartile range: 10.5 – 90). Three attending emergency physicians provided a second review for 355 patients to establish a kappa for agreement.

**Figure 1:**
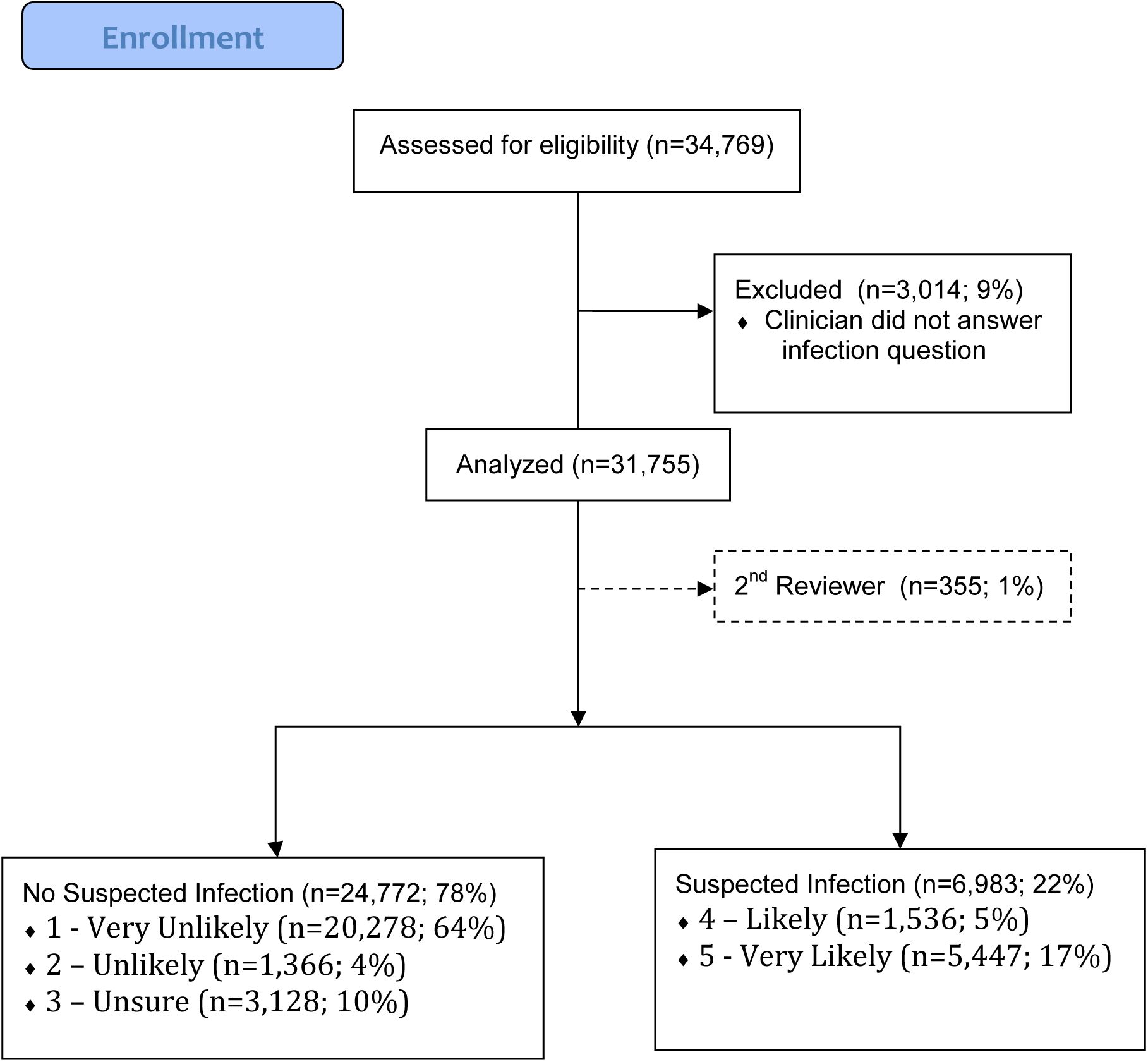
Patient Enrollment.

Patients with suspected infection (n=6,983; 22%) were slightly older (52.9 vs. 50.6), admitted more frequently (51% vs 32%; p < 0.001), admitted to the ICU more frequently 8.9% vs 5.7%; p < 0.001), spent a similar median time in the ICU of 2 days (p=0.01), spent more time in the hospital (median of 3 vs 2 days; p < 0.001), and had a higher mortality rate (1.6% vs 0.9%; p < 0.001). (**Table 1**). The original Angus sepsis abstraction criteria had a sensitivity of 55%, specificity of 97%, PPV of 82%, NPV of 88%, accuracy of 87%, and a F_1_ score of 0.66. The modified Angus sepsis abstraction criteria had a sensitivity of 65%, specificity of 96%, PPV of 81%, NPV of 91%, accuracy of 89%, and F_1_ score of 0.72. (**Table 2)**. The modified Angus criteria were more sensitive in identifying cases of suspected infection compared to the original Angus criteria (sensitivity 65% vs. 55%). The additional ICD9-CM codes caused a minimal decrease in specificity compared to the original criteria (97% vs. 96%). Overall, the modified Angus criteria outperformed the original Angus criteria as reflected by the increase in F_1_ score from 0.66 to 0.72. Overall, there was good inter-rater reliability among the primary and secondary reviewers (n=355; 1%) with a Cohen’s Kappa statistic of 0.78 (95% CI: 0.70, 0.86).

We also report a confusion matrix in **Table 3** listing the ICD9-CM codes most commonly associated with False Negatives and False Positives when using the modified Angus sepsis abstraction criteria. The ICD9-CM codes most commonly associated with false negatives were the codes for diarrhea, nausea and vomiting, abdominal pain, noninfectious diarrhea, and dehydration. The ICD9-CM codes most commonly associated with false positives were the codes for UTI, fever, acute copd exacerbation, pneumonia, and acute URI.

## Limitations

There are several potential limitations to this study. First, we used clinician assessment of suspicion of infection at time of ED disposition as the gold standard which may have misclassified patients. Patients may have been suspected of having an infection in the ED and ultimately may have had an alternative diagnosis, or vice-versa. Although we could defined our outcome measure based on formal chart review of proven infection, the American College of Chest Physicians/Society of Critical Care Medicine diagnostic criteria (6) requires only suspected infection for a diagnosis of sepsis. We also only evaluated one component of the Angus sepsis abstraction criteria. It is possible that performance characteristics would have improved if we had identified patients with both suspected infection and acute organ dysfunction. Additionally, this study occurred at a single institution and might not generalize to other institutions. Lastly, the outcome measure suspected infection was made a voluntary question in our disposition module in order to maximize the quality of collected data. It has been our experience that when clinicians are pressed for time, they often will answer mandatory questions incorrectly. Voluntary questions allow users to skip the question rather than provide spurious data. Despite voluntary reporting, we were still able to achieve a data collection rate of 91%.

## Discussion

The International Statistical Classification of Diseases and Related Health Problems (ICD) is a medical ontology designed to facilitate epidemiological surveillance. Its adaptation in the United States, the ICD9-CM, is routinely collected as part of any medical visit for coding and billing purposes. Since this information is collected for every visit, these codes are frequently used to perform epidemiological surveillance. However, the validity of this approach has been called into question when identifying ischemic cerebrovascular disease (7-11), bacteremia(12), pneumonia or acute respiratory illness(13-16), gastrointestinal bleeding(17), major and minor procedures (surgical or otherwise)(18), cardiovascular and stroke risk factors (19), healthcare-associated infection(20, 21), traumatic brain injury(22), perioperative complications(23, 24), allergies(25), diabetes(26), chronic obstructive pulmonary disease(27), Wegener’s granulomatosis(28), Clostridium difficile infection(29), and methicillin-resistant S. Aureus(30).

ICD9-CM based abstraction criteria are also widely used in epidemiological surveillance of sepsis, however its use has not been well validated. In fact, there is wide variability in the overall incidence of severe sepsis depending on how administrative data is abstracted. In the United States, Angus et al reported that the annual incidence of severe sepsis in 1995 was 3.0 per 1000(1), but Martin et al reported its incidence as 0.81 per 1000 in 2000(31, 32). By contrast, a Swedish study gives a yearly incidence of as low as 0.15 per 1000 in 1999 (33). In neighboring Norway and Finland, the incidence was reported as 0.49 per 1000 (34) and 0.38 per 1000 (35), respectively. The large difference in the incidence of severe sepsis is likely not be explained by differing demographics alone.

Wilhelms et al investigated this difference in incidence of severe sepsis by applying the three different ICD9-CM abstraction methods described by Angus et al(1), Flaatten(34), and Martin et al (31) to the same Swedish national population from 1987 to 2005. Wilhelms et al reported that these three different methodologies produced 3 entirely different populations of 37,990, 27,655, and 12,512 patients, respectively (36).

The goal of this study was to prospectively validate the accuracy of the infection component of the most commonly used sepsis abstraction method, the Angus sepsis abstraction criteria. To our knowledge, this is the first prospective clinical validation of any sepsis abstraction criteria. We found that the Angus criteria had high specificity (97%), but low sensitivity (55%). Sensitivity improved from 55% to 65% when accounting for additional ICD9-CM codes created after the original Angus publication without significant decrease in specificity (97% to 96%). These results confirm previous studies that ICD9-CM based abstraction criteria are not perfect. Furthermore, they reflect an inherent bias in emergency physician documentation and subsequent ICD9-CM code assignment. Since ICD9-CM codes are assigned based on the ED discharge diagnosis and physician documentation, these criteria tend to underestimate infection when there is not a clearly defined diagnosis. For example, ICD9-CM codes associated with Diarrhea, Nausea and Vomiting, and Abdominal Pain were the most common codes associated with False Negatives. This reflects the common clinical teaching within Emergency Medicine to document symptoms rather than diseases when the diagnosis is uncertain as a cognitive strategy to prevent diagnostic anchoring.

We tested the hypothesis that adding ICD9-CM codes for symptoms associated with infection might improve test performance. We performed a sensitivity analysis adding two ICD9-CM codes associated with False Negatives (Diarrhea, Nausea and Vomiting). Although this increased sensitivity from 65% to 72%, it reduced specificity from 96% to 91%. This reduced the F_1_ score from 0.72 to 0.71, a statistical measure that takes into account the tradeoff between sensitivity and specificity. Although these symptoms are often associated with infection, they can also be associated with non-infectious causes like chemotherapy, pregnancy, and intoxication. We did however include the symptom based ICD9-CM code for fever a priori in our modified abstraction criteria because it is specific for infection. Although there are non-infectious causes of hyperpyrexia, they are assigned different ICD9-CM codes for non-infectious causes of fever than the codes used for infectious causes of fever. We also performed a sensitivity analysis removing fever from our modified abstraction criteria, which reduced sensitivity from 65% to 59% with no change in specificity, resulting in an overall decrease in the F_1_ score from 0.72 to 0.69. The results of this sensitivity analysis are reported in **Table 4**.

Our next steps would be to prospectively validate the second component of the Angus sepsis abstraction criteria to identify acute organ dysfunction. We could then validate the combination of these two components to identify severe sepsis.

## Conclusion

In our study, the Angus abstraction criteria have high specificity (97%), but moderate sensitivity (55%) in identifying patients with suspected infection as defined by physician at the time of disposition from the emergency department. Sensitivity improved (55% to 65%) without significant decrease in specificity (97% to 96%) when additional ICD9-CM codes created after the original Angus publication were added. This resulted in an increase in the F_1_ score from 0.66 to 0.72. Given these findings, it is likely that we are underestimating the true incidence of sepsis in the United States and worldwide.

## Acknowledgements

This work was supported by CIMIT Award No. 12-1262 under U.S. Army Medical Research Acquisition Activity Cooperative Agreement W81XWH-09-2-0001. The information contained herein does not necessarily reflect the position or policy of the Government, and no official endorsement should be inferred.

SH and NIS conceived the study. All authors designed the study. SH collected the data and performed the statistical analysis. SH, JWJ, and NIS wrote the paper, and all authors contributed substantially to its revision. SH and NIS take responsibility for the paper as a whole.

